# Indigenous gut microbes modulate neural cell state and neurodegenerative disease susceptibility

**DOI:** 10.1101/2025.02.17.638718

**Authors:** Lisa Blackmer-Raynolds, Maureen M. Sampson, Anna Kozlov, Aimee Yang, Lyndsey Lipson, Adam M. Hamilton, Sean D. Kelly, Pankaj Chopra, Jianjun Chang, Steven A. Sloan, Timothy R. Sampson

**Affiliations:** Department of Cell Biology, Emory University School of Medicine, Atlanta GA 30322; Department of Human Genetics, Emory University School of Medicine, Atlanta GA 30322

## Abstract

The native microbiome influences a plethora of host processes, including neurological function. However, its impacts on diverse brain cell types remains poorly understood. Here, we performed single nucleus RNA sequencing on hippocampi from wildtype, germ-free mice and reveal the microbiome-dependent transcriptional landscape across all major neural cell types. We found conserved impacts on key adaptive immune and neurodegenerative transcriptional pathways, underscoring the microbiome’s contributions to disease-relevant processes. Mono-colonization with select indigenous microbes identified species-specific effects on the transcriptional state of brain myeloid cells. Colonization by *Escherichia coli* induced a distinct adaptive immune and neurogenerative disease-associated cell state, suggesting increased disease susceptibility. Indeed, *E. coli* exposure in the 5xFAD mouse model resulted in exacerbated cognitive decline and amyloid pathology, demonstrating its sufficiency to worsen Alzheimer’s disease-relevant outcomes. Together, these results emphasize the broad, species-specific, microbiome-dependent consequences on neurological transcriptional state and highlight the capacity of specific microbes to modulate disease susceptibility.

**Highlights:**

- The microbiome impacts the transcriptional landscape of all major brain cell types.
- Discrete microbes specifically modulate resident myeloid cell status.
- Gut *E. coli* triggers dynamic transcriptional responses across neural cell types.
- Exposure to *E. coli* exacerbates behavioral and cellular pathologies in 5xFAD mice.

## INTRODUCTION

The human body is colonized by a diverse and complex community of microbes, the microbiome, that shapes a range of host physiological processes. An individual’s gastrointestinal (GI) tract harbors 100-500 unique bacterial species that are influenced by genetic, environmental, and lifestyle factors, with an estimated 3,500 unique strains of human gut-resident bacterial species worldwide^1–4^. Given this variability in microbiome composition between individuals, across the lifespan, and within the context of disease, understanding the physiological consequences of specific microbial taxa on health outcomes, particularly neurological disease, is essential.

Gnotobiotic mouse models raised in a sterile, germ-free (GF) environment, or treated with high dose antibiotics have provided significant insights into the contributions of the gut microbiome to host physiology. For instance, in the absence of indigenous microbes, GF mice have smaller, poorly developed lymphoid organs; limited lymphocyte maturation; and increased susceptibility to infectious disease^5,6^. In the brain, GF or antibiotic-treated mice harbor immature microglia that are less capable of mounting inflammatory responses and defending against pathogens^7–9^. In addition to—or perhaps even as a result of—microbiome contributions to neuroinflammatory tone, the microbiome also has broad impacts on neurological function in the context of both health and disease. A native microbiome is necessary for proper neurogenesis, myelination, and neurotransmitter production; with impacts on anxiety-like, social, and cognitive behaviors in mice^10^. While some microbiome-dependent effects on immune and neurological functions are developmental and irreversible, others are quickly restored by microbial colonization or exposure to microbial metabolites, highlighting the importance of continuous microbial input for proper neurological functions into adulthood ^5,8,11^.

Whole microbiome manipulations in mouse models of disease have significant impacts on both behavioral and pathological outcomes^12,13^. In the absence of an intact, native microbiome, mouse models of Alzheimer’s disease (AD) show significant improvements in cognitive performance and reductions in both amyloid beta (Aß) and tau pathology^14–21^. Depletion of myeloid cells from the brains of antibiotic-treated APPPS1-21 mice eliminates the protective effect of antibiotic treatment, suggesting that microbiome-mediated neuroimmune signaling contributes to pathological outcomes^20^. While these studies emphasize the microbiome’s capacity to modulate overall brain health and function, how the microbiome shapes the transcriptional state across different brain-resident cell populations, remains poorly understood. Furthermore, despite the various associations of specific bacterial taxa with neurological diseases^12^, few studies have addressed physiological contributions of individual microbial species to relevant neuroimmune functions.

Here, we provide a single-cell characterization of microbiome-dependent transcriptional state within the mouse brain and identify both the conserved and cell-type specific transcriptional landscapes temporally modulated by microbiome-derived signals. In particular, the neuroimmune compartment was one of the most transcriptionally responsive, suggesting that these cells act to propagate signals from the microbiome to other cells within the CNS. These responses are specific, as we find that select, non-pathogenic, gut-resident species are sufficient to uniquely shape the transcriptional state of brain-resident myeloid cells. In particular, the species *Escherichia coli* induces a distinct, and temporally modulated transcriptional profile across not only brain-resident immune cells, but also across neuronal and other glial populations. Colonization with *E. coli* dynamically regulates neurodegenerative disease associated pathways across numerous neural cell types. In fact, oral exposure to *E. coli* worsens cognitive impairments and increases amyloid pathology within the 5xFAD mouse model of AD, underscoring the pathological relevance of the microbiome-dependent transcriptional landscape. Together these results highlight the widespread impact of native microbiome-dependent signaling on the brain, as well as the specific consequences of individual gut microbes for neurological function, emphasizing the importance of the native microbiome in shaping transcriptional tone that can impact disease.

## RESULTS

### A complex microbiome is necessary for the steady-state transcriptional landscape across many brain-resident cell types

In order to understand how select gut bacterial species impact the brain, we first sought to determine how the complete absence of an indigenous microbiome influenced the cellular transcriptional landscape. We performed single-nucleus RNA sequencing (snRNAseq) on hippocampal tissues derived from young adult female, germ-free (GF) or conventional (CONV) mice with a complete and intact microbiome. Unsupervised clustering of 23,963 total nuclei (representing 4 mice per treatment group) revealed seven unique clusters, roughly equally represented across each microbiome status (Fig. 1A), that were identified as the following cell types based on marker genes^22^: excitatory neurons (Cluster 1), inhibitory neurons (Cluster 2), astrocytes (Cluster 3), myelinating oligodendrocytes (Cluster 4), vasculature (Cluster 5), oligodendrocyte progenitor cells (OPCs; Cluster 6), and immune cells (Cluster 7) (Fig. 1B; Supplementary Data S1). Differential gene expression analysis demonstrated microbiome dependent transcriptional responses within each cluster, with most differentially expressed genes (DEGs) occurring within excitatory neurons, myelinating oligodendrocytes, vasculature cells, and immune cells (Fig 1C; Supplementary Data S1). Upon normalization by cluster size, to further account for the increased power in highly abundant clusters, myelinating oligodendrocytes and immune cells were found to have the largest microbiome-dependent transcriptional response (Fig 1D. This highlights these cell types’ unique susceptibility at the interface between the CNS and periphery to indigenous microbial signals.

**Figure 1.**
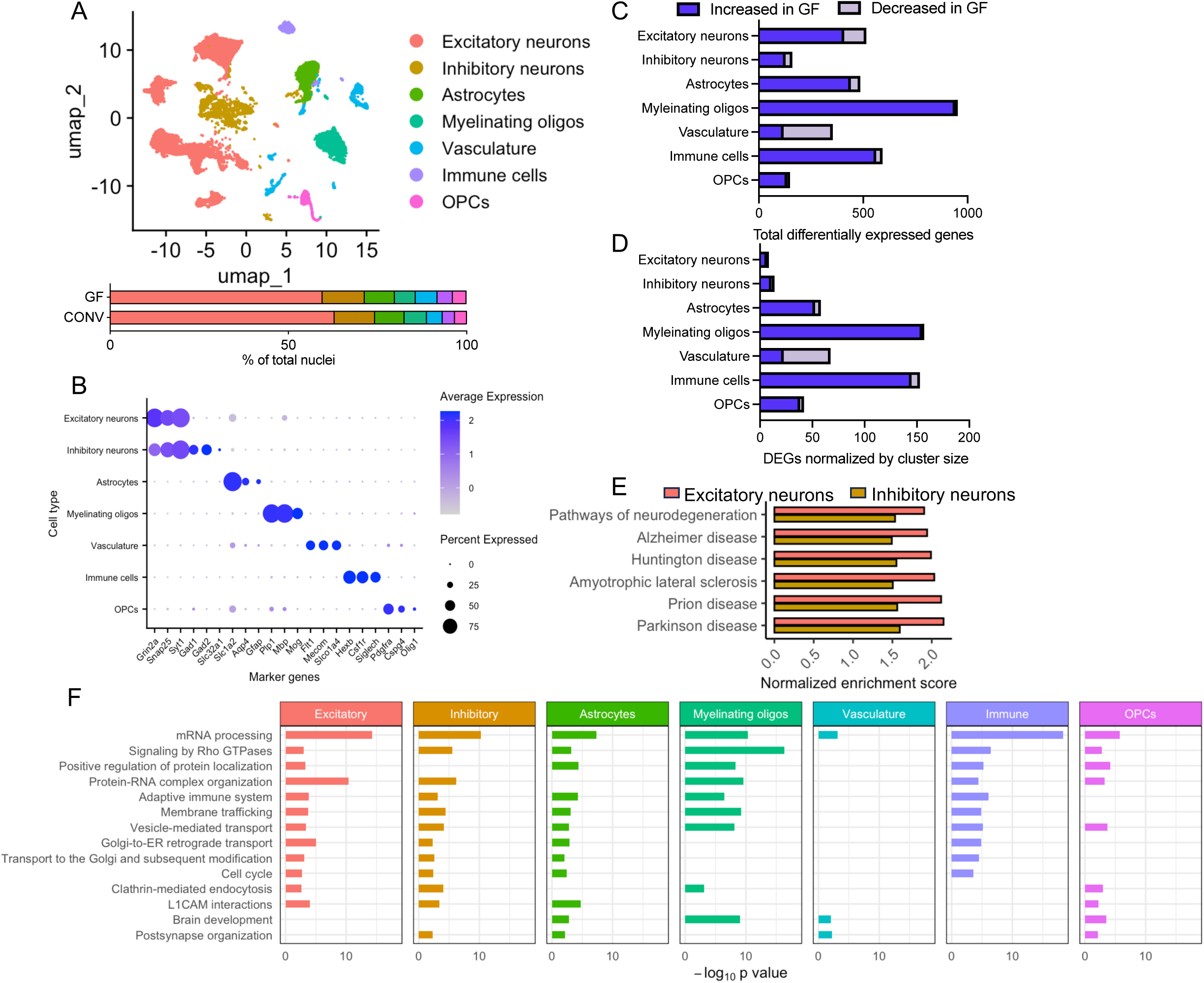
The microbiome shapes the transcriptional landscape of every major cell type in the brain **A)** Single nucleus RNA-seq was performed on hippocampal samples from female germ-free (GF) and conventionally raised (CONV) mice. UMAP shows all major cell types identified as well as the amount of each cell type found within each treatment group. **B)** Cell type markers representing each major cell type cluster. **C)** Number of differentially expressed genes (DEGs; log fold change > |1|, p < 0.001) in GF mice compared to CONV across each cell type. **D)** Gene set enrichment analysis showing increased neurogenerative disease KEGG pathways in GF mice. **E)** Representative pathways increased in at least four cell types by overrepresentation analysis (Metascape). Data represents cells from 4 mice per treatment.

To evaluate the biological relevance of these microbiome-dependent DEGs, we performed pathway analysis (Fig. 1E, F; Supplementary Fig. S1; Supplementary Data S1). We first examined biological pathways conserved in their microbiome-dependent responses across at least 4 cell types. While we did not observe any pathway decreased within 4 cell types, we did find a number of shared, microbiome-dependent features within those biological pathways enriched in many cell types (Fig. 1F). Among all cell clusters, mRNA processing (GO:0006397) was highly enriched in the absence of a microbiome. Across 4 to 5 cell types, we observed a further enrichment in protein localization (GO:0010954) and membrane trafficking (R-MMU-199991) pathways, adaptive immune system (R-MMU-1280218), cell cycle (R-MMU-1640170) and brain development (GO:0007420) pathways, indicating a widespread repression of these particular signaling pathways by the native microbiome. These observations align with prior, targeted studies which demonstrated microbiome-dependent impacts on brain development and organization^10^ and neuroimmune activation^7–9^. Even within the pathways that were increased in less than 4 cell types, impacted pathways were conceptually related, falling broadly within the categories of development, cellular stress, and immune processes (Supplementary Fig. S1A). Within both excitatory and inhibitory neuron clusters, gene set enrichment analysis (GSEA) of KEGG pathways revealed a significant microbiome-dependent enrichment of all major neurodegeneration-associated KEGG pathways (Fig. 1E). These same cell types also displayed increased immune related pathways (*e.g.* adaptive immune system (R-MMU-1280218), neutrophil degranulation (R-MMU-6798695), cellular response to interlukin-4 (GO:0070670), and interferon signaling (R-MMU-913531), among others), emphasizing the interaction between the indigenous microbiome, neuroimmune activation, and neurodegenerative disease pathways in shaping the neuronal transcriptional landscape (Supplementary Fig. S1A).

### Select gut microbes differentially and specifically modulate the brain-resident myeloid cell transcriptome

Immune cells within the brain are observed to be particularly susceptible to perturbations within the gut microbiome^7–9^ (Fig. 1D). The increases we found in immune related pathways across brain cell types, suggest that immune cells are critical for transducing microbiome-derived signaling to other cells within the brain.

However, it is unclear whether these activities are a generalized response to broad bacterial organisms or if specific, indigenous microbes are sufficient to impart differential effects on brain-resident immune cells.

Therefore, we set out to test the specificity and sufficiency of select, indigenous gut bacterial species to modulate the transcriptional state of brain-resident myeloid cells (primarily microglia). Wild-type GF mice were mono-colonized with select bacterial type strains representing prevalent genera within the mammalian gut microbiome-*Bacteroides thetaiotaomicron* (*B. theta*)*, Clostridium celatum, Lactobacillus johnsonii,* and *Escherichia coli*-for 2 weeks. CD11b+ myeloid cells, largely representing microglia but also present on other leukocytes, were then enriched^23^, and bulk transcriptomics was performed (Fig. 2A).

**Figure 2.**
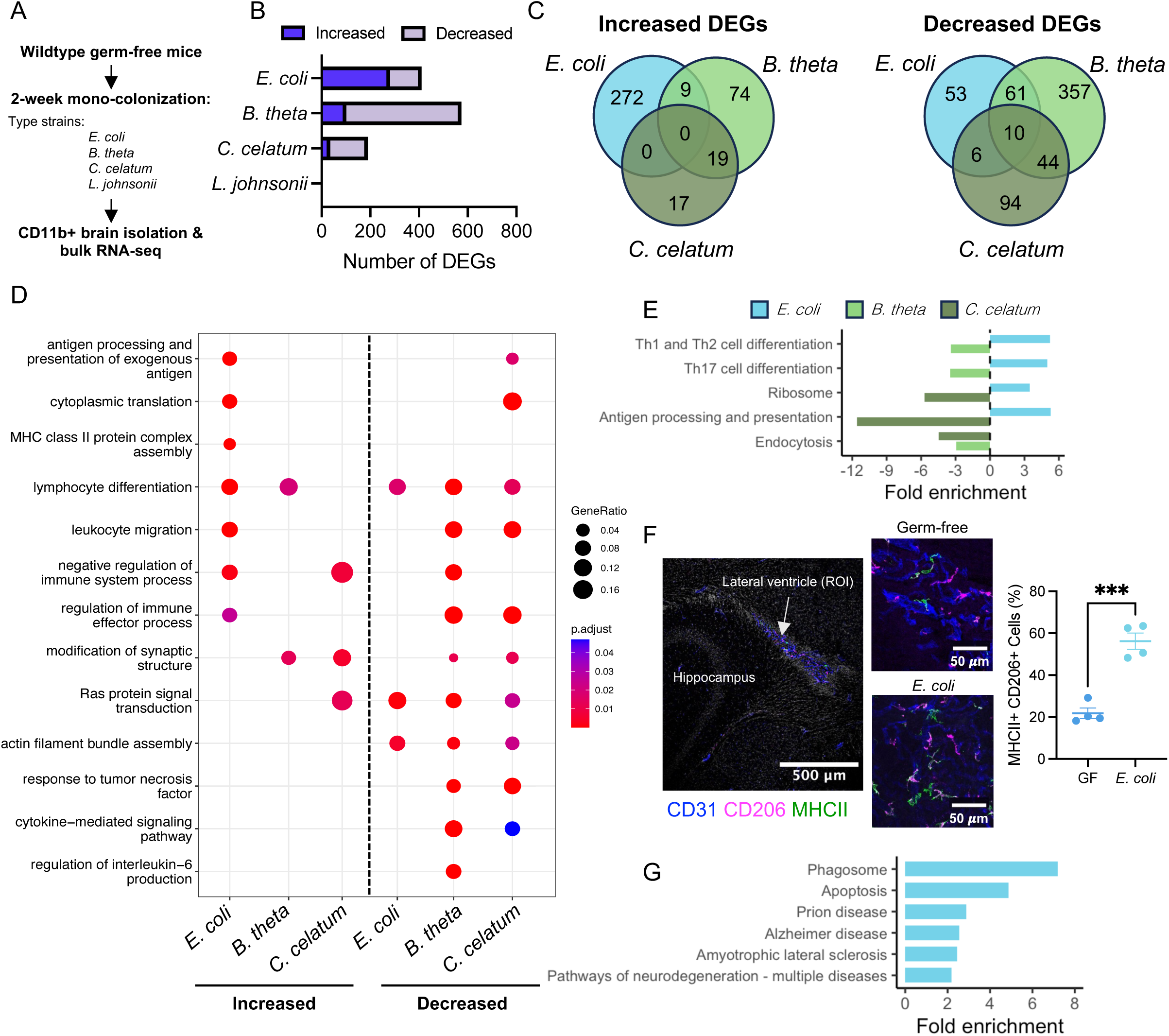
Select gut bacteria uniquely shape neuroinflammatory tone **A)** Wildtype germ-free (GF) mice were mono-colonized with type strains of interest for 2 weeks. After mono-colonization, CD11b+ myeloid cells were isolated from the brains and bulk RNA-seq was performed. **B)** Number of differentially expressed genes (log fold change > |0.25|; p < 0.05) in mono-colonized mice compared to GF. **C)** Number of overlapping differentially expressed genes across colonization states. **D)** Comparison of overrepresentation-based pathway analysis results across mono-colonization states. **E)** KEGG pathway analysis was run on both increased and decreased DEGs using DAVID. KEGG pathways significantly increased or decreased in at least two treatments are shown. **F)** Immunohistochemistry was performed on the choroid plexus of the lateral ventricle to quantify the percent of MHCII+ CD206+ double-positive CD206+ cells. **G)** KEGG pathways increased after *E. coli* mono-colonization. n = 3-7 mice. In **F)** dots represent individual mice and error bars represent SEM. *** p < 0.001.

Differential gene expression analysis of CD11b+ brain myeloid cells demonstrated that all but one of the bacteria studied—*L. johnsonii*—were sufficient to modulate myeloid cell gene expression, with colonization by *E. coli* and *B. theta* inducing the most DEGs (Fig. 2B; Supplementary Data S2). Comparison of the DEGs increased and decreased after mono-colonization shows very little overlapping DEGs, highlighting the specificity of the transcriptional response to each of these unique bacterial species (Fig. 2C). No shared DEGs were increased by all three species and of the decreased genes, only 10 were shared across colonization states (Fig. 2C). Further emphasizing the unique transcriptional state induced by each bacterial species, pathway analysis comparing across each bacterial colonization state shows near inverse effects between the unique species (Fig. 2D; Supplementary Data S2). Where *E. coli* induced an increase in pathways involved in immune activation and adaptive immune responses, *B. theta* and *C. celatum* decreased many of these same pathways in male mice. Further emphasizing the unique transcriptional response of brain myeloid cells to *E. coli*, comparison of KEGG pathway enrichment revealed that *E. coli* mono-colonization triggered an inverse effect to that of *B. theta* or *C. celatum* amongst all shared KEGG pathways (Fig. 2E). To validate the transcriptional increase in genes involved in adaptive immune processes, specifically MHCII antigen presentation, immunohistochemistry was performed to evaluate MHCII levels within the brain of male mono-colonized mice (Fig. 2F). MHCII+ cells were primarily found within CD206+ cells, canonically considered to be border associated macrophages, within the CD31+ choroid plexus of the lateral ventricle. In line with the transcriptional data, when compared to GF mice, *E. coli* mono-colonization induced significantly more MHCII+ / CD206+ double-positive cells (Fig. 2F), supporting the increase in adaptive immune transcriptional processes within the brains of these animals. In addition to pathways involved in adaptive immune responses, *E. coli* alone induced an increase in KEGG pathways involved in neurodegenerative diseases—including prion disease (mmu05020), AD (mmu05010), and Amyotrophic lateral sclerosis (ALS, map05014)—highlighting the potential for *E. coli* colonization to modulate a disease-susceptible transcriptional state within the brain (Fig. 2G). Notably, multiplex measurement of cytokines in both intestinal tissues and serum demonstrated limited impacts to inflammatory cytokine levels (Supplementary Figure S2A, B). While a decrease in ileal TNF and increase in colonic CXCL1 and TNF was observed in most mono-colonized conditions (Supplementary Fig. S2A), we observed no overt effect in colon length or other measure of GI function (Supplementary Fig. S2C-E). In conjunction with no observed effect on circulating cytokines during colonization by any tested microbe (Supplementary Fig. S2B), these data suggest that mono-colonization did not induce robust local or systemic inflammatory signaling. Similarly, no overt gliosis was observed, with limited impact on the IBA1+ area (Supplementary Fig. S2F, G), demonstrating transcriptional effects in the absence of robust gliosis or systemic inflammation.

### *E. coli* elicits a temporal transcriptional response across CNS-resident cells

Given the experimental contributions of *E. coli* to pathologies in models of neurodegenerative disease^24–27^, and our observations here that this native microbe is sufficient to drive both adaptive immune and neurodegenerative disease-associated pathways in brain-resident myeloid cells, we next sought to understand how this organism dynamically shapes transcriptional tone across many neural cell types. We performed snRNA-seq on hippocampal samples from female mice mono-colonized with *E. coli* for either 2 or 4 weeks to capture both short- and longer-term transcriptional responses following microbial colonization in comparison to GF controls. Clustering of 30,093 total nuclei (representing 4 mice per treatment group) was performed in combination with previously assessed nuclei (Fig. 1) to create comparable cell clusters (Fig. 3A, B; Supplementary Data S3). Interestingly, while cell numbers between each cluster appeared relatively unchanged following 2 weeks of *E. coli* colonization, by 4 weeks we observed a qualitative shift in cell populations. This is highlighted by over an 18.5% decrease in excitatory neuron representation and a corresponding increased representation of inhibitory neurons, myelinating oligodendrocytes, and astrocytes (Fig. 3A).

**Figure 3.**
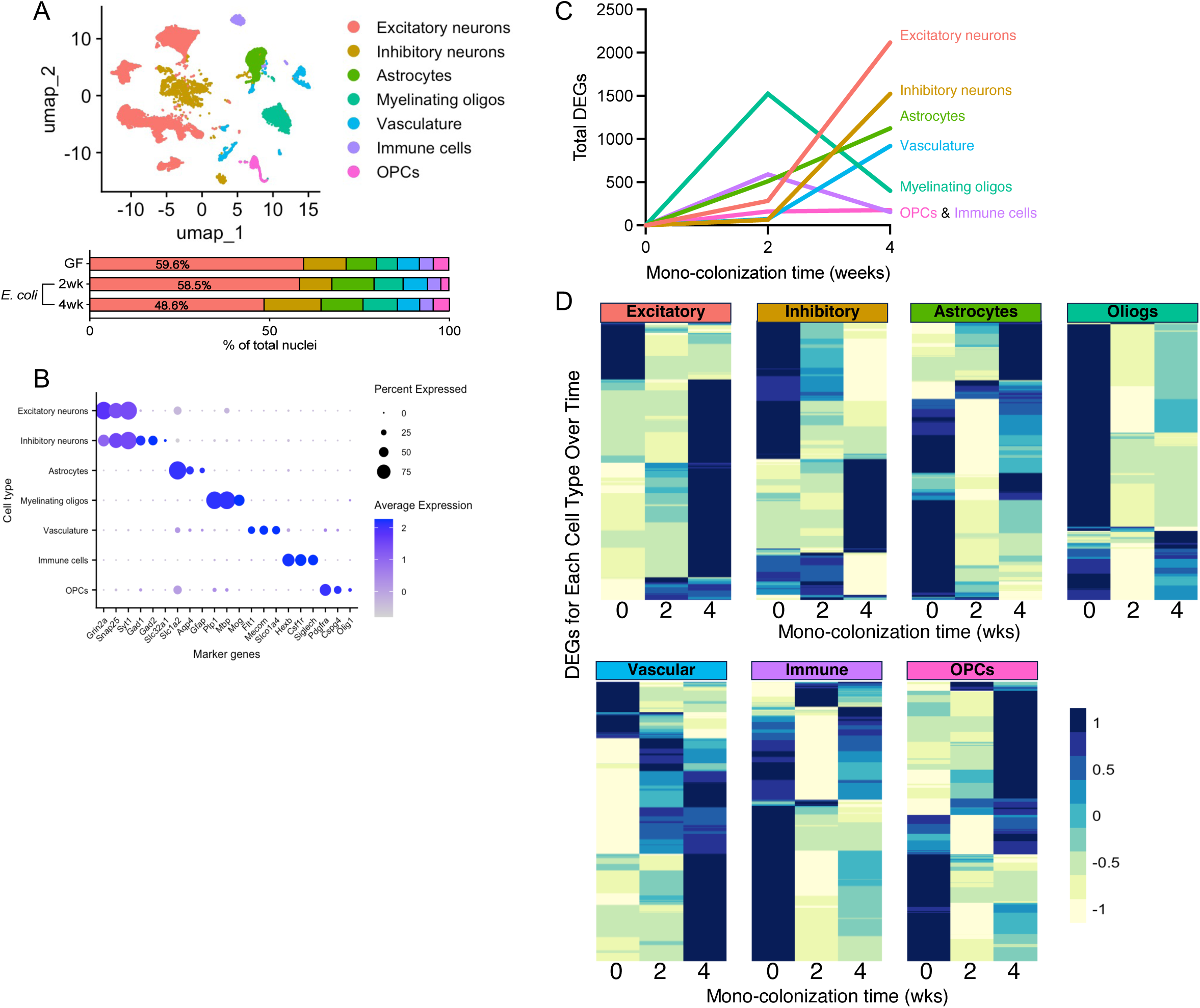
Gut colonization with *E. coli* temporally modulates gene expression across all cell types the brain Single nucleus RNA sequencing was performed on hippocampal samples from germ-free (GF) mice and mice that were mono-colonized with *E. coli* for either 2 or 4 weeks. **A)** UMAP showing cell clusters and bar graph showing the percentage of total nuclei within each cluster based on treatment. **B)** Cell type markers for each major cell type cluster. **C)** Graph showing the total number of differentially expressed genes (DEGs; log fold change > |1|, p < 0.001) compared to GF per cell type across time. **D)** Heat map shows relative expression levels of all differentially expressed genes (at either 2 or 4 weeks) within each cell type across time (note each heat map represents only that cell type’s DEGs). Data represents cells from 4 mice per treatment.

Differential gene expression analysis showed time dependent shifts in the number of DEGs increased in each cell type compared to GF (Fig. 3C; Supplementary Data S3). Where the immune cell cluster displayed the most increased DEGs and myelinating oligodendrocytes the most decreased DEGs at 2 weeks post-colonization, the number of DEGs in both of these cell types was dramatically reduced at 4 weeks. In contrast, the number of DEGs in the other cell clusters (excluding OPCs that remained largely unresponsive) was dramatically higher at 4 weeks compared to 2 weeks. While many of the DEGs that were significantly altered at 2 weeks in the immune cell and myelinating oligodendrocyte clusters began to stabilize back towards a non-colonized state at 4 weeks (Fig. 3D), they did not fully recover, highlighting long-term, yet subtle, transcriptional states that may alter susceptibility to future insults. In contrast, the neuron clusters—with the highest number of DEGs at 4 weeks (Fig. 3C)—displayed a limited transcriptional response until 4-weeks post-colonization with *E. coli* (Fig. 3D). We interpret this to suggest that these cells are either slower to respond to microbiome status or instead, respond to those signals derived from more acutely responsive cells.

Since we observed the immune cell cluster as having the most increased DEGs in the early colonization state at 2 weeks post-*E. coli* colonization (Fig. 3), and bulk RNA-seq of brain myeloid cells displayed a microbiome-dependent adaptive immune and neurodegenerative disease phenotype (Fig. 2), we hypothesized that this cell type may initiate the broader transcriptional responses in other cells throughout the brain. In order to distinguish the microbiome dependent effects on microglia compared to other immune cell types, we performed sub-clustering and pathway analysis within the immune cell cluster (Fig. 4; Supplementary Data S4).

**Figure 4.**
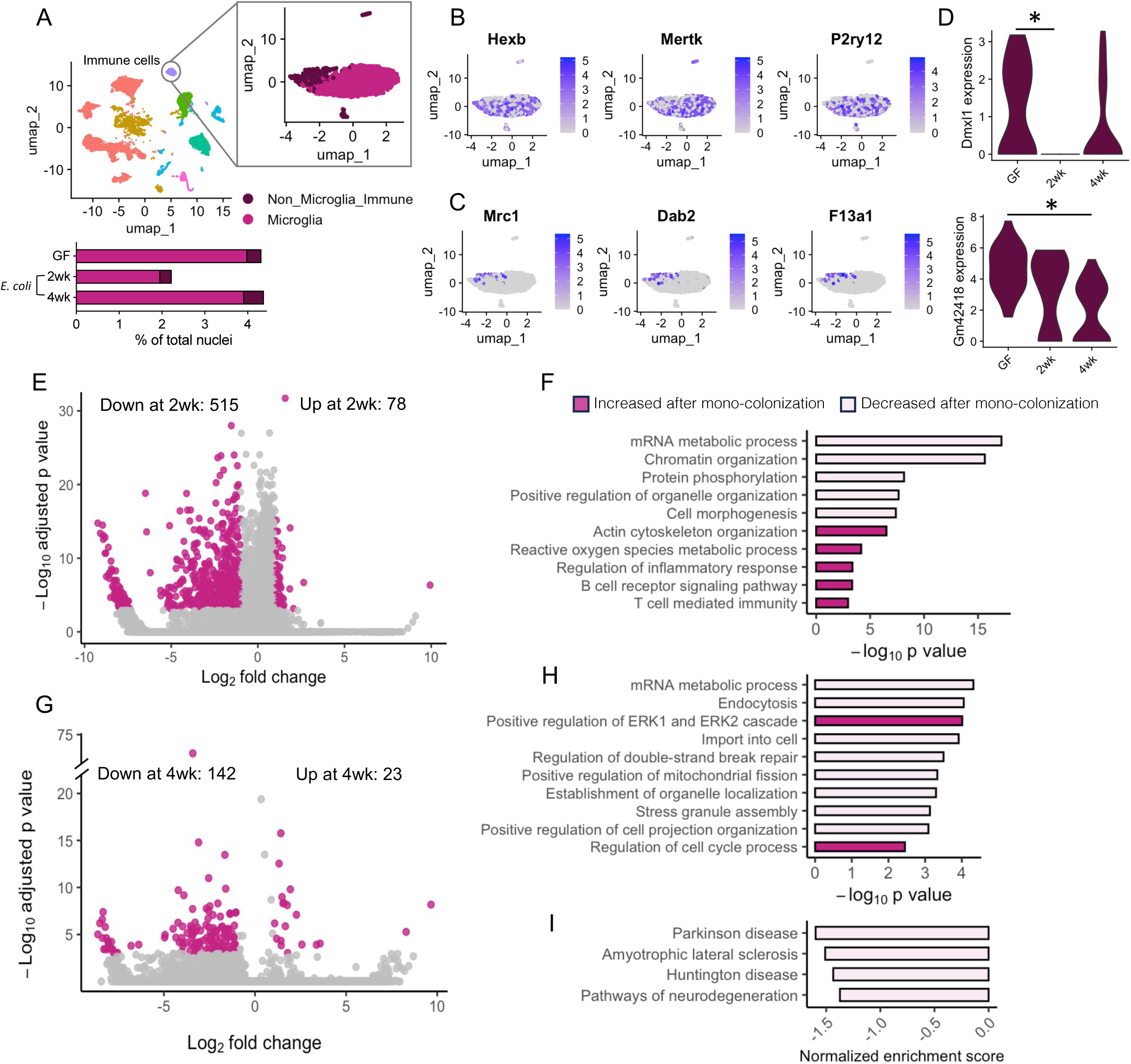
*E. coli* colonization induces temporal regulation of microglial transcriptional state. The immune cell cluster identified in Figure 2 was further subclustered to distinguish microglia from non-microglia immune cells. **A**) Shows cell clustering UMAP and % of all nuclei within each cell type. Markers defining each cell cluster are overlayed upon the immune UMAP with **B)** representing microglia markers and **C)** representing non-microglia markers. Differential gene expression analysis was performed only both cell clusters. **D)** shows violin plots of the two DEGs that reached significance in the low powered non-microglia immune cluster. **E)** Volcano plot and **F)** pathway analysis of microglia differentially expressed genes (log fold change > |1|, p < 0.001) after 2 weeks of *E. coli* mono-colonization. **G)** Volcano plot, **H)** GO pathway analysis and, **I)** neurodegeneration KEGG GSEA of microglia after 4 weeks of mono-colonization. Cells represent 4 mice per treatment.

Unsupervised sub-clustering identified 2 main sub-clusters: microglia and non-microglia immune cells (*e.g.* border associated macrophages, infiltrating peripheral immune cells) (Figs. 4A-C). While similarly prevalent irrespective of colonization status, non-microglia immune cells were lowly abundant and showed little transcriptional responsiveness to *E. coli* across both time points (Fig. 4D), however, this may be due to an insufficient sample size, rather than a lack of susceptibility to perturbation.

In contrast, differential gene expression analysis identified 515 repressed and 78 induced genes in microglia following 2-weeks of mono-colonization (Fig. 4E). Pathway analysis on these DEGs identified similar pathways as we observed in our analysis of CD11b+ cells (Fig. 2) including an increase in pathways involved in inflammatory responses and adaptive immunity, whereas downregulated pathways included those characterized as cellular organization and mRNA processing (Fig. 4F). At 4-weeks post-colonization, differential gene expression analysis of microglia revealed 23 increased and 142 decreased DEGs (Fig. 4G). We observed a repression in genes involved mRNA metabolic processes and cellular organization, as we observe at the 2-week time point. However, microglia also showed a decrease in cellular stress pathways (*e.g.* regulation of double stranded break repair (GO:2000779) and stress granule assembly (GO:0034063)) at 4-weeks post-colonization (Fig. 4H). Further, GSEA of microglia at the 4-week timepoint displayed a significant decrease in a range of neurodegenerative disease pathways (Fig. 4I), contrasting with the *E. coli* triggered increase in these pathways we observed at 2-weeks (Fig. 2G).

To better understand the breadth of the transcriptional response to *E. coli* colonization, we performed pathway analysis within every other cell cluster at each timepoint (Fig. 5A, B; Supplementary Data S5). At both 2- and 4-weeks post-colonization, we observed a significant decrease in pathways involved in RHO GTPase signaling, adaptive immunity, and RNA metabolism across cell types. This mirrors our observations in mice with a complex, and intact microbiome (Fig. 1E), and highlights that these transcriptional pathways may be highly sensitive to microbial colonization. In contrast, pathways involved in nervous system / brain development were increased in *E. coli* colonized mice, while these were decreased in conventional mice compared to GF animals (Fig. 1E). Cell-specific pathways further highlight that colonization with *E. coli* induced a time-dependent response in immune, cellular stress, and cellular organization/transport pathways (Supplementary Fig. S3).

**Figure 5.**
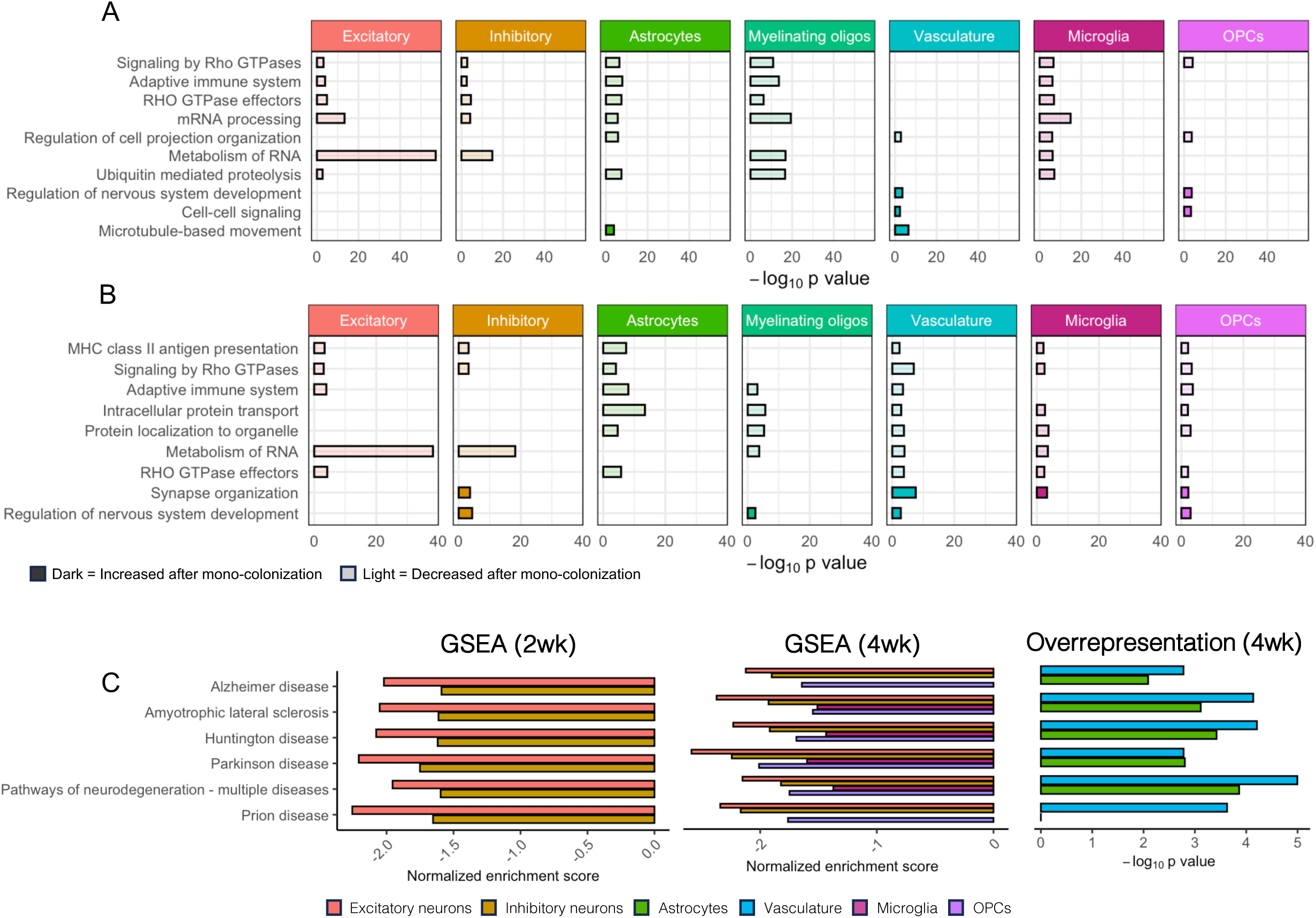
*E. coli* mono-colonization modulates biological pathways involved in adaptive immune responses and neurodegeneration across cell types and timepoints Overrepresentation pathway analysis was run using the DEGs (log fold change > |1|, p value < 0.001) between GF and *E. coli* mono-colonized mice at each timepoint. Increased and decreased pathways that overlapped the most across cell types at 2 weeks are shown in **A)** and at 4 weeks are shown in **B)**. Gene set enrichment analysis was also performed to identify neurodegeneration KEGG pathways that are modulated by *E. coli* colonization. **C)** Shows the normalized enrichment score for each cell type with a significant GSEA result at 2 and 4 weeks as well as neurodegeneration KEGG pathways increased in overrepresentation analysis at 4 weeks. Data is from 4 mice per treatment.

Since these transcriptional responses are relevant for neurological disease, we specifically examined neurodegenerative disease KEGG pathways by both overrepresentation pathway analysis and GSEA (Fig. 5C). We observed a modulation of neurodegeneration KEGG pathways across nearly every cell type following *E. coli* colonization, with a decrease in KEGG pathways of neurodegenerative diseases in both excitatory and inhibitory neurons following colonization across both timepoints, and in microglia by 4-weeks post-colonization (Fig. 5C). In addition, overrepresentation-based pathway analysis demonstrated an enrichment of neurodegeneration pathways in both astrocytes and vasculature at 4 weeks of colonization (Fig. 5C). Taken together, these results emphasize a link between *E. coli* colonization and transcriptional pathways that are involved in modulation of neurodegenerative disease risk.

### *E. coli* modulates cognitive impairment in an animal model of amyloid pathology

We have demonstrated that *E. coli* is sufficient to transcriptionally modulate adaptive immune and neurodegenerative disease pathways across many cell types in the brain. Notably, *E. coli* and closely related organisms have been reported to be enriched within the gut microbiome of individuals with neurodegenerative disease^12,28–32^, suggesting it may contribute to disease processes. We therefore sought to directly test the disease modulatory potential of *E. coli* within the context of AD—the most prevalent neurodegenerative disease. To evaluate the sufficiency of gastrointestinal *E. coli* to modulate AD outcomes, we used the well-characterized 5xFAD mouse model^33^, that displays amyloid pathology beginning at 2 months of age^34^ and cognitive impairment between 3 and 6 months of age^35^. To test exacerbation over this model’s pathological progression, 2-month-old conventionally raised 5xFAD mice were orally exposed to ∼10^8^ cfu of a non-pathogenic, strain of *E. coli* (identical to our mono-colonization experiments) or vehicle control 3x weekly for 1 month prior to behavioral and pathological assessments (Fig. 6A). Irrespective of treatment, 5xFAD mice displayed similar outcomes in both the open field and intestinal behaviors, suggesting that *E. coli* exposure did not induce overt sickness or anxiety-like behaviors, and GI functions were not robustly disrupted (Supplementary Fig. S4 A-E). While working memory, as measured in the Y-maze test, appeared intact (Fig. 6B), *E. coli* exposure resulted in a loss of novelty preference in the object location test (Fig 6C). Similarly, while learning capacity during Barnes Maze training was not impacted (Fig. 6D), *E. coli* treated 5xFAD mice displayed a significantly longer primary latency during the probe trial suggesting a loss of typical memory function (Fig. 6E). These observations appeared specific to the 5xFAD genotype, as wildtype littermates did not demonstrate any loss of cognitive functions in an identical battery of tests (Supplementary Fig. S5). Thus, in a genotype specific fashion, exposure to intestinal *E. coli* is sufficient to exacerbate cognitive decline in this mouse model.

**Figure 6.**
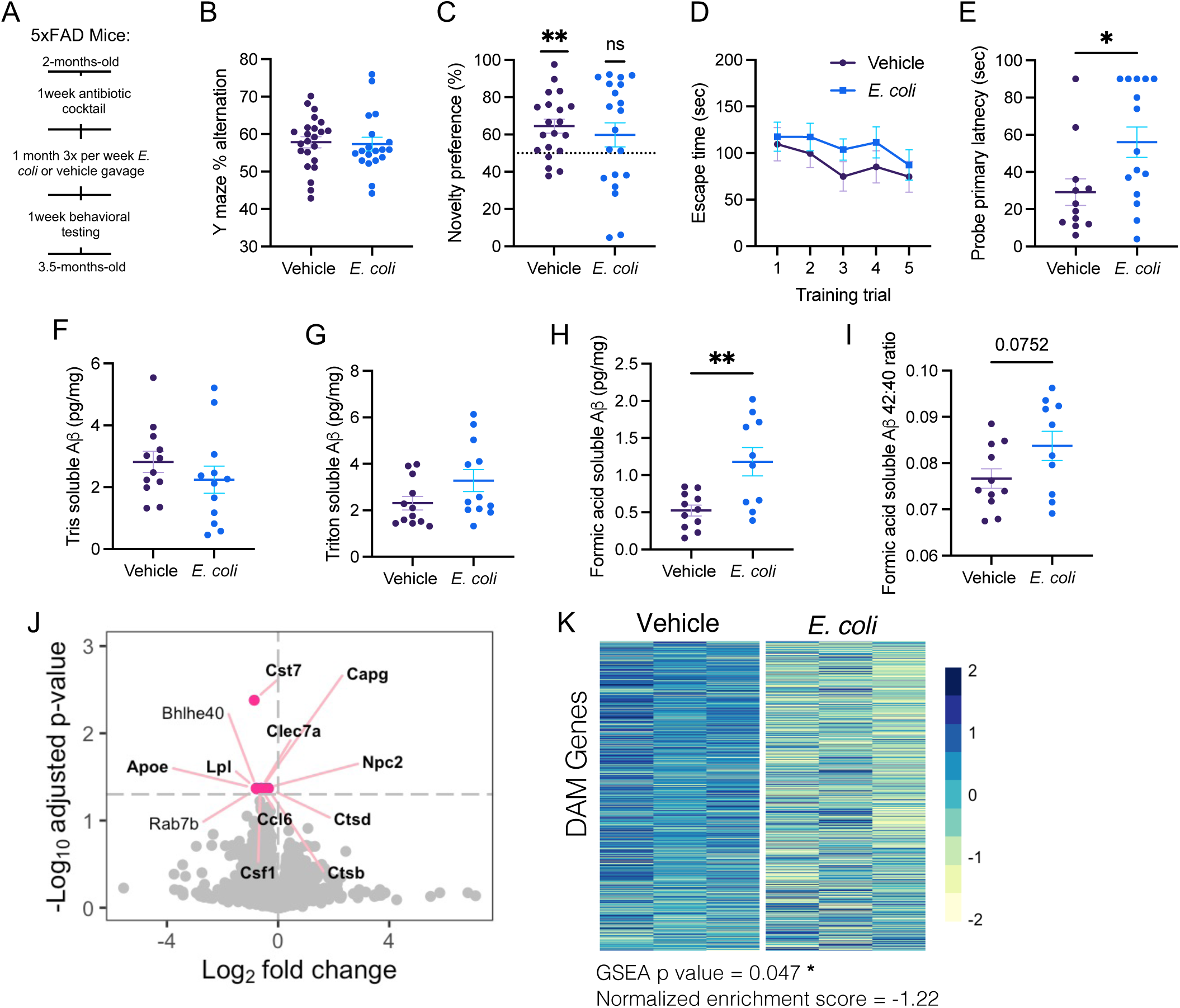
*E. coli* exposure exacerbates AD outcomes in 5xFAD mice **A)** Male and female 2-month-old 5xFAD mice underwent a 1-month enrichment paradigm followed by behavioral and pathological assessments. Behavioral testing was performed on the **B)** Y maze, **C)** Object location test, and **D-E)** Barnes maze to evaluate different forms of spatial memory function. Levels of amyloid beta were measured within **F)** tris soluble, **G)** triton soluble, and **H)** formic acid soluble protein fractions extracted from the hippocampus. **I)** The ratio of amyloid beta 42:40 as also calculated the formic acid soluble fraction. Bulk RNA-seq was performed on CD11b+ myeloid cells in the brain with **J**) showing a volcano plot of differentially expressed genes with disease associated microglia (DAM) genes highlighted in bold and **K)** showing the expression level of all DAM genes as well as results of a gene set enrichment analysis (GSEA). For **B - I** n = 11-15, dots represent individual mice, bars represent mean ± SEM. Groups were compared using a two tailed t test except **C** where each condition was compared to the 50% chance level using a one sample t test. **J-K)** n = 3 dots **(J)** and rows **(K)** represent individual genes and columns represent individual mice **(K)**. * p < 0.05 ** p < 0.01.

To evaluate whether intestinal *E. coli* exacerbates pathological outcomes associated with the cognitive impairments observed in 5xFAD mice, we measured hippocampal Aß concentrations (Fig. 6 F-I). Where there were no differences in the concentration of Tris- or triton-soluble Aß (Fig. 6F-G), *E. coli* treated animals had significantly more formic acid soluble Aß (representing Aß in a highly insoluble form) than the vehicle treated controls (Fig. 6H), along with a trend towards increased Aß 42:40 ratio (Fig. 6I). Together, these data suggest that *E. coli* exposure exacerbates insoluble amyloid deposition. Profiling of cytokines and chemokines across anatomical compartments including the colon, serum, and hippocampus, demonstrated that exposure to *E. coli* did not induce a robust pro-inflammatory response in any anatomical compartment, in line with our behavioral evaluation for sickness-like behaviors (Supplementary Fig. S4 F-H).

Bulk RNA-seq of brain-derived CD11b+ cells from 5xFAD mice demonstrated a disease relevant transcriptional response to *E. coli* exposure. Brain myeloid cells, derived from *E. coli* treated mice had 12 repressed DEGs (Fig. 6J). Despite a small number of DEGs, nearly all are highly relevant for AD. For example, Apoe—the greatest known genetic risk factor for late onset AD^36^—had one of the largest log fold change values, and all but two of the DEGs (Bhlhe40 and Rab7b) are known to be increased in the classical disease associated microglia (DAM) phenotype^37^. In support of this association, GSEA highlights a significant decrease in DAM genes following *E. coli* exposure (Fig. 6K). This suggests that *E. coli* exposure results in an inability for brain myeloid cells to transition into the initially protective DAM state and may explain the worsened cognitive behaviors and pathology observed in *E. coli* exposed 5xFAD mice. Overall, these results highlight how gut exposure to non-pathogenic *E. coli* alters brain wide transcriptional state in healthy wildtype animals and accelerates disease progression in a genetic model of AD.

## DISCUSSION

Increasing experimental evidence demonstrate that the gut microbiome maintains constant communication with the brain, shaping neurological function in both health and disease^10,38^. We find that the gut microbiome shapes the transcriptional landscape of every major cell type in the brain, highlighting the breadth of microbiome-derived signaling. At a single cell resolution, we identify both cell-type specific and conserved transcriptional responses dependent on the presence of an intact microbiome. Further, our data delineates the shared and unique transcriptional responses elicited by particular gut microbial taxa, demonstrating the capacity for specific microbes to evoke distinct responses that are relevant for health and disease. Colonization with *E. coli* induces a broad transcriptional activation state associated with adaptive immune and neurodegenerative disease pathways and further exacerbates disease outcomes in a mouse model of AD. Together the present study highlights the association between the gut microbiome community, the active transcriptional landscape in the brain, and neurological disease susceptibility.

Over the past several decades, numerous studies have underscored the importance of the gut microbiome in shaping neurological function including consequences for a wide range of neurological cell types^10,38^. In line with this, we observe a consistent pattern, shared across cell types, of microbiome-dependent influences on developmental and cellular organization processes that are not typically found within adult mouse brains.

Previous studies have demonstrated that microbiome-derived signals are necessary for appropriate neuroimmune development, with GF mice displaying an immature microglia phenotype and an inability to mount a typical inflammatory response^7–9^. Our data demonstrate a global dysregulation of genes associated with adaptive immune processes in the brain, in the absence of a microbiome, further emphasizing the importance of the microbiome for neuroimmune processes. Even in healthy, wildtype mice, both excitatory and inhibitory hippocampal neurons display dysregulated transcription of genes assigned to neurodegenerative disease pathways. These findings support the emerging role of the microbiome in modulating neurodegenerative disease outcomes, including numerous observations of microbiome-dependent pathology in both genetic and toxicant-induced models of neurodegenerative disease^12,13^.

The composition of the gut microbiome differs across individuals, including significant differences in those living with neurological conditions^4^. It is therefore important to not only understand the consequences of the microbiome as a whole, but also to pinpoint the specific effects of individual gut microbes on neurological functions. This allows a deeper understanding of whether and how particular microbial associations may ultimately contribute to neurological outcomes. Indeed, we found species-specific effects on transcriptional activation of neuroimmune cells, including one organism *L. johnsonii*, whose colonization induced no transcriptional modulation compared to GF controls. This surprising finding demonstrates that neuroimmune cells do not simply respond broadly and non-specifically to microbial colonization, but that there is indeed specificity in these interactions arising from the gut environment. Three other organisms, *B. theta, C. celatum,* and *E. coli* each induced unique transcriptional responses, demonstrating the specific neuroimmune modulatory capacity of these GI resident bacterial taxa. *E. coli* notably triggered an increase in expression of genes involved in antigen presentation and adaptive immune activation, pathways which were strikingly decreased with colonization by both *B. theta* and *C. celatum*. Similarly, colonization with *E. coli* induced expression of several pathways associated broadly with neurodegenerative diseases including prion disease, AD, and ALS within brain myeloid cells, emphasizing the disease modulatory potential of this bacterium.

Microbiome-dependent transcriptional responses in the brain are dynamic. Our temporal analysis identified that immune cells and myelinating oligodendrocytes are acutely responsive to *E. coli* colonization, while neuronal populations do not display a response until 4 weeks post-colonization. Expression of those genes involved in adaptive immune pathways were initially robustly upregulated within microglia following colonization, but this response subsided by 4 weeks post colonization. However, the increase in immune pathways, particularly those involved in adaptive immunity, became apparent in nearly every other cell type by 4 weeks post colonization, suggesting that microglia are a focal point in relaying microbiome-derived signals to other cells in the brain. This solidifies prior observations of microbiome-dependent shifts in microglia state during both development and in models of neurodegenerative disease. It further suggests that microbiome-elicited microglia responses may trigger subsequent transcriptional impacts across other cell types in the brain.

Colonization with *E. coli* was sufficient to induce a transcriptional response associated with neurodegenerative diseases in wildtype mice. We further observed that increased exposure to *E. coli* within the 5xFAD mouse model of AD accelerated the development of cognitive decline and amyloid pathology. Cognitive impairments were associated with decreased expression of genes within the classical DAM phenotype, thought to be an initially protective microglia state that limits the development and progression of AD outcomes^37,39^. This lack of protective response within brain myeloid cells may help to explain why disease outcomes progressed more rapidly in *E. coli* exposed mice. Interestingly, we and others, have observed that *E. coli* and related taxa within the *Enterobacteriaceae* family are sufficient to exacerbate pathologies in other models of neurodegeneration^24–27^. Our data herein expand on this concept, identifying specific brain transcriptional responses evoked by the presence of *E. coli* within the microbiome that may contribute to these outcomes. This is of particular importance given observations that *Enterobacteriaceae* are enriched in the gut microbiome amongst many neurodegenerative diseases^12^.

While we utilized a model of AD-relevant pathologies, our data serve as a foundation to understand how microbiome-dependent transcriptional responses associated with specific microbial species can modulate neurological disease susceptibility. Systemic immune responses vary greatly in response to individual species, even within the same genera^40^, suggesting that delineating individual microbial contributions is essential. While *E. coli* exemplifies how non-pathogenic microbiome derived signals shape the neurological transcriptional landscape, our data demonstrate that individual bacterial species—and perhaps bacterial strains—will induce differential outcomes. For example, we highlight the seemingly inverse consequences of *B. theta* on neuroimmune transcriptional state compared to *E. coli,* yet it has been demonstrated that exposure to a related species of Bacteroides, *Bacteroides fragilis,* is sufficient to exacerbate AD outcomes^41–43^. While both *Escherichia* and *Bacteroides* have been reported to be increased in people living with AD^29–32,44–47^, public studies of the AD-associated microbiome are currently limited. Understanding the species-specific associations in these human conditions and their experimental contributions to neurological functions remains an important gap in the field. Together, our results highlight the specificity and dynamics of microbiome-derived signals on the transcriptional landscape of the brain, as a foundation for the continued study of how the indigenous microbiome shapes overall brain health and disease susceptibility.

## RESOURCE AVAILABILITY

### Lead contact

Further information and requests for resources should be directed to and will be fulfilled by the lead contact, Timothy Sampson.

### Materials availability

This study did not generate new, unique materials.

### Data and code availability

Transcriptomic data is deposited into the NIH GEO database and is available upon acceptance. All original code will be publicly deposited in GitHub and will be freely available upon acceptance. DOI and accession numbers are listed in the key resources table. Any additional information required to reanalyze the data reported in this paper is available from the lead contact upon request.

## ACKNOWLEDGMENTS

We thank Isabel Fraccaroli, Hsiao-Lin Wang, and Emily Hill for scientific support; and all the members of the Emory Division of Animal Resources for technical support. We acknowledge support from the Emory Multiplexed Immunoassay Core (EMIC), the Emory Integrated Cellular Imaging Core (ICI), the Emory Integrated Genomics Core (EIGC), and the Emory Gnotobiotic Animal Core (EGAC) which are subsidized by the Emory University School of Medicine as Integrated Core Facilities and are supported by the Georgia Clinical and Translational Science Alliance of the NIH (UL1TR002378). This work is funded by NIH/NIEHS 1R01ES032440 (TRS); NIH/NIA F31AG076332 (LBR); NIH K00 ES033033 and Burroughs Welcome Fund Postdoctoral Enrichment Program (BWF PDEP) (MMS); T32 NS096050 (AMH); and NIH/NIMH R01MH125956 (SAS). The content is solely the responsibility of the authors and does not necessarily reflect the official views of the sponsors.

## AUTHOR CONTRIBUTIONS

LBR and TRS conceived and designed the study. LBR, AK, AM, LDL, AMH, SDK, and JC performed the experiments. MMS, PC, and SAS provided computational analysis. LBR analyzed the data and prepared figures. TRS supervised the study. LBR and TRS wrote the manuscript. All authors revised and approved the manuscript.

## DECLARATION OF GENERATIVE AI AND AI-ASSISTED TECHNOLOGIES IN THE WRITING PROCESS

During the preparation of this work, the authors did not use generative AI or AI-assisted technologies.

## STAR+METHODS

### EXPERIMENTAL MODEL AND STUDY PARTICIPANT DETAILS

#### Animals

##### Gnotobiotics

Germ-free (GF), male and female DBA/2N mice were originally obtained from Taconic Biosciences (#DBA2; RRID: IMSR_TAC:DBA2) following embryonic rederivation and bred within the Emory Gnotobiotic Animal Core (EGAC), for at least 3 generations prior to use in this study. GF animals were co-housed (with 2-5 same sex and age matched cage mates) in sterile cages within Parkbio rigid isolators. Mice were provided sterile food (Teklad Autoclavable Diet, 2019S) and water *ad libitum*. Microbiological testing (by culture and qPCR) was performed on all autoclaved materials entering isolators (including food, water, and bedding) as well as monthly within the isolators themselves. Prior to sacrifice or mono-colonization, all mice were transferred to sterile, static housing in specific pathogen-free (SPF) vivarium. Mono-colonization was subsequently performed by oral gavage with ∼10^8^ cfu of bacteria of interest within a sterile, class II biological safety cabinet. Colonization status of all mice was confirmed by fecal culture at the time of sacrifice.

##### Conventionally-reared mice

Conventionally-raised (CONV) male and female DBA/2J mice were originally obtained from Jackson Laboratory (#000671; RRID: IMSR_JAX:000671) and co-housed (2-5 per cage) in static housing with food (LabDiet: 5001) and water provided *ad libitum* within an SPF facility. Female and male 5xFAD mice, on a congenic C57Bl/6J background (Jackson Labs, #034848; MMRRC_034848-JAX) were maintained by crossing with C57Bl/6J wildtype mice (IMSR_JAX:000664). Mice were co-housed (2–5 per cage) with mixed genotype littermates in sterile, microisolator cages, under a 12h light/dark cycle with *ad libitum* access to sterile food (Teklad Autoclavable Diet, 2019S) and drinking water. Genotypes were confirmed with the following vendor-approved primers and their PCR parameters: APP Forward 5′-AGGACTGACCACTCGACCAG-3′, APP Reverse 5’-CGGGGGTCTAGTTCTGCAT-3′; PS1 Forward 5′-AATAGAGAACGGCAGGAGCA-3′, PS1 Reverse 5′-GCCATGAGGGCACTAATCAT-3′. All animal husbandry and experiments were performed in accordance with AVMA guidelines and approved by the Institutional Animal Care and Use Committee of Emory University (PROTO201900056).

##### Bacteria

*Bacteroides thetaiotaomicron* str. VPI 5482 (ATCC 29148), *Clostridium celatum* str. VPI 8759-1 (ATCC 27791), and *Lactobacillus johnsonii* str. VPI 7960 (ATCC 33200) were obtained from the American Type Culture Collection (ATCC). *Escherichia coli* str. K12 MC4100 (a kind gift from Matthew Chapman (University of Michigan^24^) and *B. thetaiotaomicron* were grown aerobically at 37° C in tryptic soy broth (BD #211825) and brain heart infusion (BD #237500) supplemented with hemin and vitamin K, respectively. *C. celatum* and *L. johnsonii* were grown anaerobically at 37° C in Chopped Meat Carbohydrate (Anaerobe Systems AS-811) and de Man-Rogosa Sharpe (BD 288130) broth respectively. Bacterial cultures were resuspended at ∼10^8^ cfu in sterile 50% glycerol and 5% sodium bicarbonate, plated to confirm monoculture, and stored at −80° C until use. Vehicle control consisted of sterile 50% glycerol (v/v) and 5% sodium bicarbonate (w/v) also stored at −80° C.

## METHOD DETAILS

### Bacterial enrichment

Male and female, conventionally raised 2-month-old 5xFAD and wildtype littermates were treated with an antibiotic cocktail consisting of 1mg/mL neomycin, 1mg/mL ampicillin, and 5mg/mL vancomycin in sterile water for 1 week. After antibiotic treatment, mice were randomly assigned to receive either ∼10^8^ cfu of *E. coli* by oral gavage 3x per week or vehicle (sterile 50% glycerol and 5% sodium bicarbonate) gavage. While cages consisted of mixed genotype animals, each cage of mice only received a single treatment to prevent cross-contamination. Each week, the colonization status of the mice was monitored by fecal culture.

### Behavioral Testing

Roughly equal numbers of male and female 5xFAD (n = 11-15) and wildtype (n= 9-23) littermates underwent behavioral testing after 1 month of enrichment, at 3 months of age. All behavioral testing was performed during the animal’s light cycle. Before the start of any test, mice were habituated to the testing room in their home cage for 1hr. All behavioral tracking and analysis were performed using EthoVision XT software (Noldus Information Technology, Wageningen, the Netherlands) and the testing arenas/objects were cleaned between trials with 70% ethanol to eliminate bacterial contamination and olfactory cues. Mice were tested on the following tests in order:

### Open field test (OFT)

As measures of motor and anxiety-like behavior, mice were placed in a 45cm square open field box for 10 minutes and allowed to explore freely. Distance traveled and time spent in the center (20×20cm) was recorded. The OFT also served as habituation for the object location test (OLT).

### Object location test (OLT)

Twenty four hours after the OFT, the OLT was run to assess short term spatial memory as described step-by-step in^61^. The same open field box used in the OFT was used for the OLT, but additional landmarks (large papers with stripes, stars, or dots of different colors) were placed on 3 out of the four walls to allow for spatial orientation. During the initial study phase, mice were placed in the box with two identical copies of an object (either a plastic chess piece or 5mL Eppendorf tube) and allowed to explore freely for 10 minutes. The mice were then returned to their home cage for a 10-minute retention delay. During the testing phase, mice were returned to the box where one of the two objects had been moved to a novel location and allowed to explore freely for 5 minutes. Object exploration was considered time spent with the mouse’s nose within 2cm of an object. Novelty preference was measured by taking the percent of total object exploration time that was spent exploring the moved object. A novelty preference significantly above 50% chance levels is indicative of intact memory as mice generally seek out the moved object.

### Y-maze

In order to evaluate spatial working memory capacity, the Y maze test was performed as described in detail in^62^. Mice were placed in a plastic Y shaped maze and allowed to explore freely for 8 minutes while the order of entries into each arm of the maze was recorded. Percent alternation was calculated by taking total number of alternations (consecutive entries into 3 arms before repeating any arms) divided by the maximum possible alternations (total entries minus 2) and multiplying by 100. Mice with intact working memory display higher percent alternation as they prefer to explore previously unvisited areas.

### Barnes maze

Longer term spatial learning and memory was evaluated using the Barnes maze test adapted from^63^, and described in detailed in^64^. Testing was performed over a 6-day period using a 92cm diameter, 20-hole, Barnes maze (MazeEngineers) with consistent extra-maze cues throughout. First, mice underwent a habituation trial in which they were placed on the maze with bright lights and white noise (66-70 dB) playing for 20 seconds then gently guided to an escape hole leading to a dark box filled with sterile bedding. After entering the escape box, the white noise was immediately turned off and the mice were allowed to acclimate to the escape box for 2 minutes. The mice then underwent 5 consecutive training trails spread over 2 days. During each training trial, mice were placed in the center of the maze (with white noise and bright lights on) and allowed to explore for up to 3 minutes. If the mice entered the escape box during this period, the noise was turned off and they were allowed to rest for 1 minute. Mice that did not enter the escape box after 3 minutes were gently guided to the proper hole before being allowed to rest. The probe trial occurred 72hr after completion of the final training trail. During the probe trial, the escape box was removed, and mice were placed on the maze (lights and noise on) to explore for 90 seconds. During both training and probe trials, primary latency (the time it took for the mouse to first check the escape hole) was recorded by hand.

### Intestinal behaviors

To evaluate whether colonization disrupts gastrointestinal (GI) function, a subset of roughly equal male and female mono-colonized (n=6-15) and 5xFAD mice (n=5-9) underwent a set of GI tests. First, colonic transit was measured by fecal output testing described in detail in^65^. Fecal output testing was performed within a level II biosafety cabinet within the animal’s vivarium with no prior habituation period. Mice were placed in individual sterile 1L plastic beakers for 30 minutes and the number of fecal pellets produced was recorded every 5 minutes. Fecal water content was measured by collecting the fecal pellets produced during the fecal output test and weighing them before and after drying at 100°C for 48hr. As a measure of total intestinal transit time, carmine red dye elution was performed as described in^66^. Testing was performed in the behavioral testing room. Mice were allowed to acclimate for 1hr before undergoing oral gavage with 100µl of sterile carmine red dye (6% w/v) (Sigma, C1022) dissolved in 0.5% methylcellulose (w/v; Sigma, M7027). Mice remained in their home cages for 2 hours then were transferred to individual sterile cages devoid of bedding, but with access to a small amount of sterile food and water. The cages were observed every 15 minutes for the presence of a red fecal pellet. Once a red pellet was discovered, the time was recorded, and the mouse was returned to its home cage.

### Tissue collection and processing

Mice were humanely euthanized via open-drop isoflurane overdose followed by cardiac puncture to collect blood samples, perfusion with PBS, and exsanguination. To collect serum, blood was immediately placed into vacuette collection tubes (Griener #454243) and spun at room temperature at 1,800 g for 10 minutes. Serum was then stored at −80° C until use. Brain tissue was either A) immediately processed by immunopanning or magnetic activated cell sorting (MACS; see below); B) dissected and flash frozen in liquid nitrogen and stored at −80° C for later protein or nuclei isolation; or C) fixed for 24hr in 4% paraformaldehyde and stored at 4° C in sodium azide for immunohistochemistry. Intestinal tissue was removed from the mouse and colon length was measured as a marker of overall intestinal inflammation. Approximately 1cm of tissue was dissected from the ilium and proximal colon, flash frozen in liquid nitrogen and stored at −80° C for multiplex ELISA.

### Brain myeloid cell enrichment

Immediately after sacrifice, brains were homogenized using a Wheaton dounce tissue grinder (with 0.114 ±0.025mm pestle clearance) in culture media comprised of HBBS with 1.5% HEPES, and 0.5% glucose. Cells were then transferred to a 35% isotonic percoll gradient solution and spun 20 minutes at 700 x g to remove the myelin layer. Transcription (Actinomycin D, Sigma #A1410) and translation (Anisomycin, Sigma #A9789) inhibitors were added to every solution throughout both procedures to prevent transcriptional changes that could impact downstream results. The remaining cells were washed several times in HBBS culture media before undergoing either immunopanning or magnetic activated cell sorting (MACS) separation.

### Immunopanning

Cells were resuspended in 0.02% BSA in PBS and incubated at room temperature on a 10cm petri dish with anti-CD11b and secondary antibody already adhered to the dish (Thermo Scientific, 14-0112-82 and Jackson 112-005-167) for 10-15 minutes. After incubation the unadhered cells were washed away and remaining cells scraped off the plate directly into Trizol (Zymo Research R2050-1-200) and stored at −80° C until later use.

### MACS

Sorting was performed according to the manufacturers (Miltenyi Biotec) guidelines. Samples were resuspended in PB buffer (0.5% BSA in PBS pH 7.2) with CD11b MicroBeads (Miltenyi Biotec # 130-093-634) and incubated for 15 minutes at 4° C protected from light. Cells were then washed and resuspended in PB buffer before being placed on an LS column (Miltenyi Biotec #130-042-401). Magnetic separation was repeated on a second LS column to maximize purity. After separation, cells were resuspended in Trizol and stored at −80° C for later use.

### Single nuclei preparation and sequencing

Flash frozen hippocampal samples from 12–15-week-old female germ-free, conventionally raised, and mono-colonized mice (n = 4) underwent a nuclei isolation protocol adapted from^67^. Frozen tissue was placed in homogenization buffer (consisting of 0.26M sucrose, 0.03M KCl, 0.01M MgCl_2_, 0.02M Tricine-KOH pH 7.8, 0.001M DTT, 0.5mM Spermidine, 0.05mM Spermine, 0.3% NP40, and protease inhibitor) and homogenized using both pestle A (0.0030-0.0050 in. clearance) and pestle B (0.0005-0.0025 in. clearance) of a KIMBLE dounce tissue grinder (Sigma D8938). Density gradient centrifugation was performed using iodixanol and the nuclei band was captured at the 30%-40% iodixanol interface. Nuclei were washed in RSB buffer (0.01M Tris-HCL pH 7.5, 0.01M NaCl, 0.003M MgCl_2_, and 0.1% Tween-20) then samples were combined such that each treatment group was represented by two samples, each containing roughly equal numbers of cells from two mice within a single treatment group. Then, samples immediately underwent cell capture using the Chromium Next GEM Single Cell 3’ Kit v3.1 (10x Genomics # PN-1000268) according to the manufacturer’s guidelines. Approximately 1,600 cells were loaded onto a Chromium Next GEM Chip and run using a Chromium Controller and library preparation steps were performed according to the manufacturer’s guidelines. Pooled samples were sequenced on a NovaSeq X Plus 25B 2×150 for 5-6.25B PE reads total by Admera Health (South Plainfield, NJ).

### Single nucleus RNA-seq analysis

Reads were aligned to the mouse reference genome (GRCm39) using the Cell Ranger pipeline (version 4.0.10, 10x Genomics). The Cellbender^50^ (version 0.3.0) remove-background function was used to minimize the effects of ambient RNA (expected cells=5,500 and total droplets included=12,000). The snRNA-seq libraries were imported into R (version 4.3.2) using Seurat^51^ (version 5.1.0) then filtered to only include cells with 200-7,500 features and less than 6% mitochondrial reads. Data was normalized, scaled, and clustered using Seurat defaults and doublets were removed using DoubletFinder^52^ (version 2.0.4) with a multiplex rate of 0.8% per 1,000 cells. Samples where integrated using Seurat RPCA Integration then re-clustered to create an integrated UMAP. Seurat “FindAllMarkers” function was used (min.pct = 0.25, thresh.use = 0.25) to identify marker genes for each cluster. Manual cell type annotation was performed by identifying cell types with high marker gene expression within the Allen Brain Cell Atlas^22^. Differential expression analysis was performed within each cell cluster of interest by using MAST within the Seurat “FindMarkers” function. Genes were considered differentially expressed if they had an absolute log fold change value greater than 1 and adjusted p value of less than 0.001. Functional annotation clustering was performed using Metascape^53^ (version 3.5.20240901) with all DEGs except genes without a canonical name (those ending in “rik” or beginning with “Gm”) and GSEA was run using clusterProfiler^59^ (version 4.10.1).

### Bulk RNA-seq preparation and sequencing

RNA was extracted from isolated brain myeloid cells using the Qiagen RNeasy Kit (#74106) according to the manufacturer’s guidelines. Bulk RNA-sequencing libraries were created for each sample using the Takara SMART-Seq mRNA HT LP kit (#634792) according to the manufacturer’s protocol and recommendations. Sequencing was performed by Admera Health (South Plainfield, NJ) using a NovaSeq or HiSeq to achieve approximately 85 million total reads per sample.

### Bulk RNA-seq analysis

Sequencing quality control was performed using FastQC (version 0.11.9) and reads were pseudoaligned to the mouse reference genome (GRCm39) using Kallisto (version 0.44.0)^54^. Files were imported into R (version 4.3.2) and counts were converted into TMM normalized log_2_ counts per million. Genes expressed in less than 3 samples were excluded from the analysis. Differential gene expression analysis was performed using limma^55^ (version 3.58.1) and edgeR^56^ (version 4.0.16). Genes were filtered to only include genes enriched in microglia compared to other brain cell types based on expression levels published by the Brain RNA-seq Website^48^.

Genes were considered differentially expressed if they had a Bonferroni adjusted p value less than 0.05. Functional annotation clustering was performed using DAVID^57,58^ and clusterProfiler^59^ (version 4.10.1), gene set enrichment analysis was performed by fgsea^60^ (version 1.28.0).

### Protein extraction and ELISA

Protein was extracted from flash frozen brain and gut samples by sonication in homogenization buffer consisting of 125mM Tris, 15mM MgCl_2_, 2.5mM EDTA (PH 7.2), 1% Triton X 100, and protease inhibitor (Roche #11697498001, 1 tablet per 10mL). Samples were centrifuged at 20,800 rcf for 10 minutes at 4° C. The protein concentration of the supernatant was measured using a Pierce BCA Protein Assay Kit (Thermo #23225) according to the manufacture’s guidelines. When quantifying amyloid beta levels, a three part protein solubility protocol outlined in^68^ was performed instead. First, the tris soluble fraction was isolated by sonication in 125mM Tris, 15mM MgCl_2_, 2.5mM EDTA (PH 7.2), and protease inhibitor (Roche #11697498001, 1 tablet per 10mL). Following centrifugation, the pellet was then re-sonicated in the above buffer with the addition of 1% Triton X 100 to isolate the triton soluble fraction. Finally, the remaining pellet was sonicated in buffer containing 70% formic acid to extract the most insoluble protein fraction. Protein samples as well as pure serum samples were then run using multiplex ELISA (Meso Scale Discovery V-PLEX Proinflammatory Panel 1 (mouse) Kit # K15048D and V-PLEX human amyloid beta peptide kit #K15200E).

### Immunohistochemistry

Paraformaldehyde fixed brain hemispheres were transferred to a 30% sucrose solution for 24-48hr then frozen in Tissue-Tek O.C.T. compound (#4583) and sliced into 40μm coronal sections using a cryostat. Two to four sections representing the anterior hippocampus including the 3^rd^ and lateral ventricles were stained per mouse. Antigen retrieval was performed prior to IBA1 staining by heating the tissue to 90° C for 5 minutes in a sodium citrate antigen retrieval buffer (10 mM tri-sodium citrate dihydrate and 0.43mM Tween 20, pH 6.0). Tissue was then blocked in 1% BSA, incubated in anti-IBA1 antibody (Wako #019-19741 1:1,000), and anti-rabbit 594 secondary (Thermo #A11012, 1:1,000). Imaging was performed using a Keyence BZ-X Series microscope (Itasca, IL) at 20X magnification. Images were processed in Fiji using a macro created by the Emory Integrated Cellular Imaging Core that auto thresholded the images using “RenyiEntropy dark,” converted to mask, and calculated IBA1% area within a hand traced region of interest around the hippocampus. MHCII, CD31, and CD206 staining was performed without antigen retrieval using the following antibodies and concentrations:

MHCII (BioLegend #107602) 1:300 with anti-rat biotin (Thermo #A187843) and streptavidin 488 (Thermo #S31354); CD31(R&D Systems # AF3628) 1:400 with anti-rabbit 594 (Thermo #A21207) 1:1,000; and CD206 (Cell Signaling # 24595T) 1:400 with anti-goat 647 (Thermo #A21447). A streptavidin/biotin blocking kit (Vector labs SP-2002) was used in conjunction with biotinylated antibodies to improve MHCII signal. Images were taken at 20x using a Leica SP8 multiphoton microscope and analyzed using Fiji “Li” and “Moments” auto thresholding for MHCII and CD206 respectively. Microscopy and image analysis was performed by a blinded lab member.

## QUANTIFICATION AND STATISTICAL ANALYSIS

### Overview of Statistical Tests

Unless otherwise indicated in the figure legends, data are expressed as mean ± SEM. Sample sizes are indicated in the figure legends, and when possible, each individual sample is represented as its own point on each graph. In all analyses except snRNA-seq, a sample represents an individual mouse. In SnRNA-seq a single sample represents combined data from two mice of the same treatment. Statistical tests for all non-transcriptomic data were performed using GraphPad Prism 8. One-way ANOVAs were used to compare mono-colonized mice and T tests were used to compare enriched (5xFAD and wildtype) mice. The object location test was analyzed by a one sample T test comparing each treatment to the 50% chance level. Unless otherwise noted significance was determined to be a *p* value of less than 0.05. Details on transcriptomic data analysis can be found in their respective sections above. All unique code is publicly available on GitHub upon acceptance.

