## Supplementary Info for "Indigenous gut microbes modulate neural cell state and neurodegenerative disease susceptibility"

**Supplementary Figure S1**, *associated with Fig. 1*. Pathways altered in GF mice by cell type.
**Supplementary Figure S2**, *associated with Fig. 2*. Neurodegeneration-associated gut bacteria modulate intestinal, but not circulating cytokine levels or microgliosis.
**Supplementary Figure S3**, *associated with Fig. 5*. *E.* coli modulates unique biological pathways for each cell type at 2 and 4 weeks.
**Supplementary Figure S4**, *associated with Fig. 6*. *E. coli* exposure does not induce sickness behavior or robust inflammation in 5xFAD mice.
**Supplementary Figure S5**, *associated with Fig. 6*. *E. coli* exposure does not induce cognitive impairment in wild type mice.

**Supplementary Figure S1.**

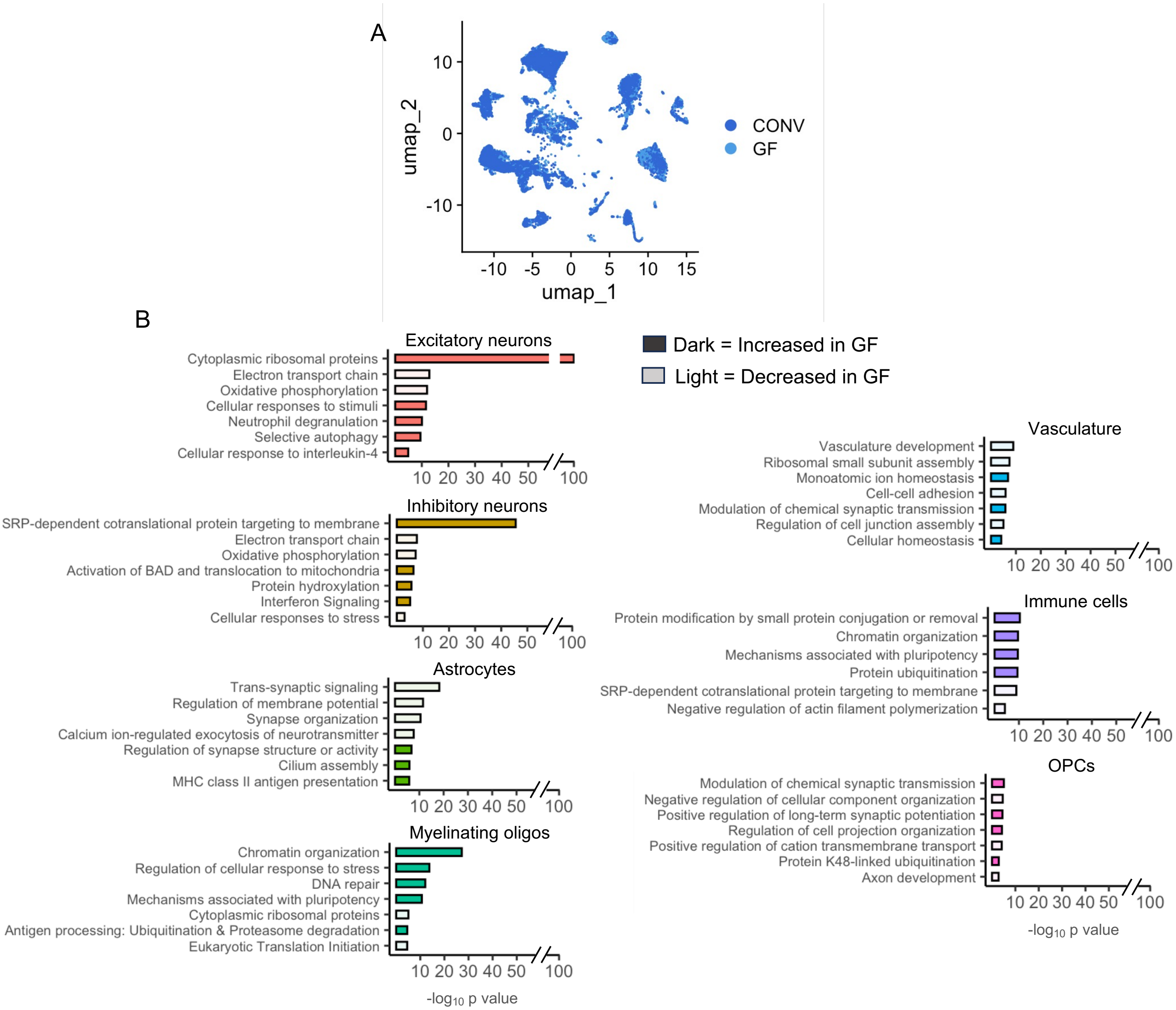


**Supplementary Figure S1, *associated with Fig. 1*. Pathways altered in GF mice by cell type. A)** Hippocampal single-nucleus RNAseq, as in Fig. 1, UMAP with nuclei highlighted by colonization status. **B)**Overrepresentation based pathway analysis was run on every major cell cluster using Metascape (log fold change > |1|, p < 0.001). Representative pathways that are increased and decreased in each cell type (and not included in Fig. 1E) are shown. Cells are from 4 mice per treatment group.

**Supplementary Figure S2.**

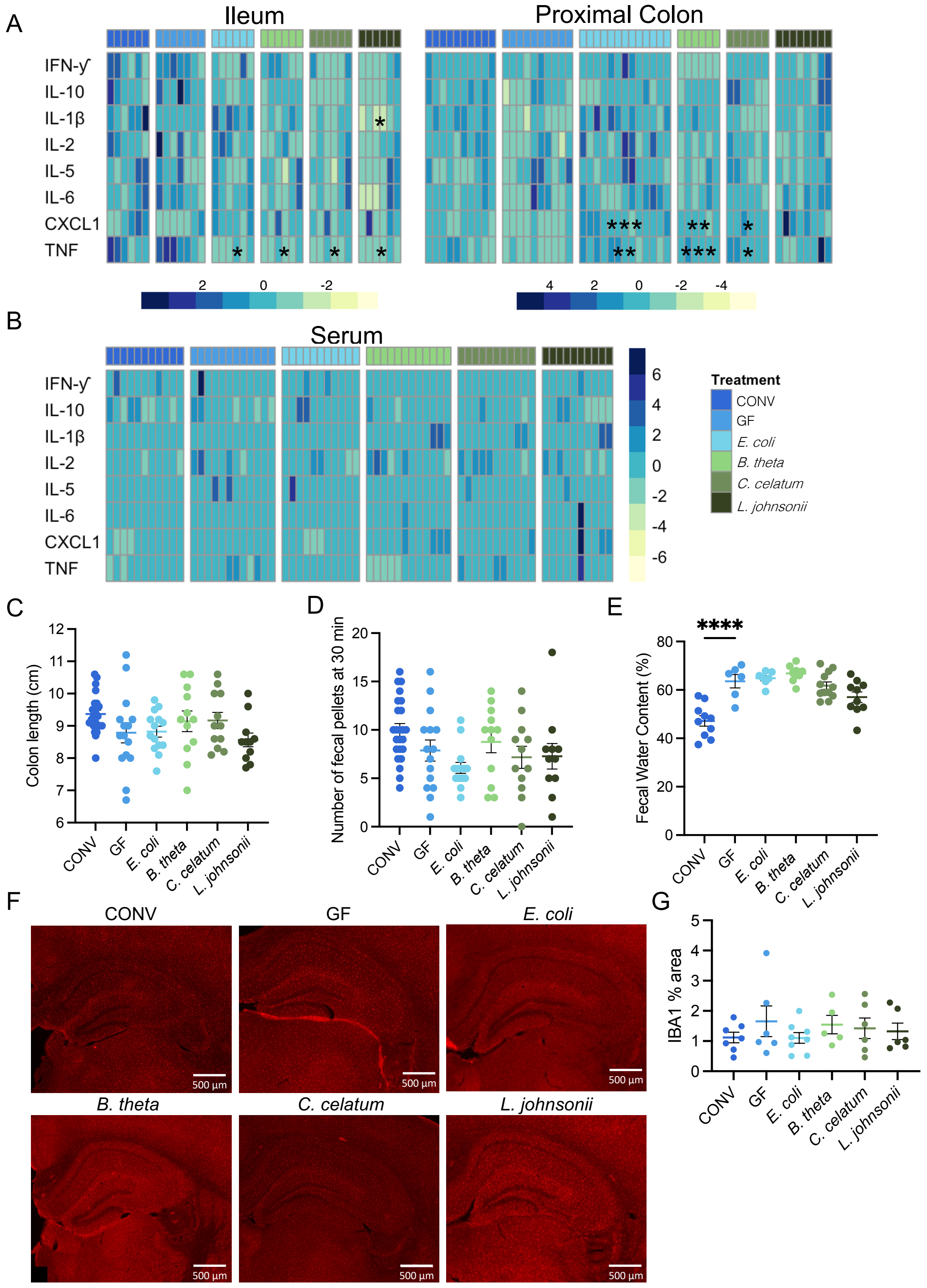


**Supplementary Figure S2, *associated with Fig. 2*. Neurodegeneration-associated gut bacteria modulate intestinal, but not circulating cytokine levels or microgliosis. A)**. Inflammatory cytokines and chemokines were measured in the ileum, proximal colon, and **B)** serum by multiplex ELISA. **C)** Colon length at the time of sacrifice was recorded as a measure of generalized colonic inflammation. **D)** Fecal output and **E)** fecal water content was used to assess gastrointestinal function. **F)** IBA1 staining was performed in the hippocampus and **G)** % area was compared between groups. n = 5-24. Dots (graphs) or columns (heat maps) represent individual mice. Error bars represent mean ± SEM. All treatment groups were compared to germ-free (GF) using either two-way repeated measures ANOVAs (**A-B**) or one-way ANOVAs (**C-E, G**) with Dunnett's multiple comparison tests**.** * $p < 0.05$; ** $p < 0.01$; *** $p < 0.001$; **** $p < 0.0001$ compared to GF. Conventionally colonized (CONV).

### Supplementary Figure S3.

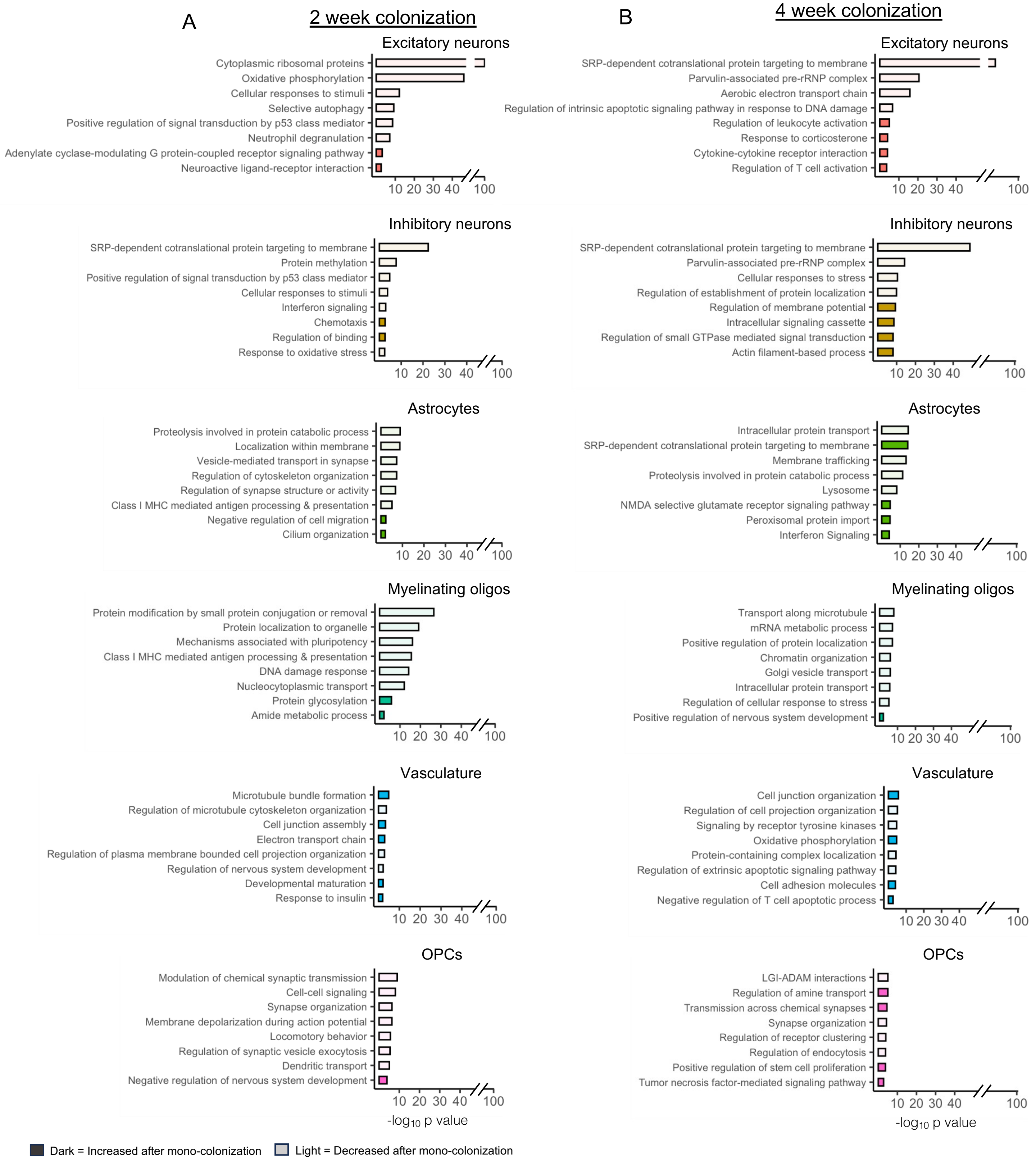


**Supplementary Figure S3, *associated with Fig. 5. E.* coli modulates unique biological pathways for each cell type at 2 and 4 weeks.** Overrepresentation based pathway analysis was run on the increased and decreased DEGs for each cell type (log fold change > |1| , p <0.001) after 2 and 4 weeks of mono-colonization using Metascape. Representative pathways (excluding those shown in Figure 5) for the 2-week timepoint are shown in **A)** and 4-week timepoint are shown in **B).** Data is from 4 mice per treatment.

### Supplementary Figure S4.

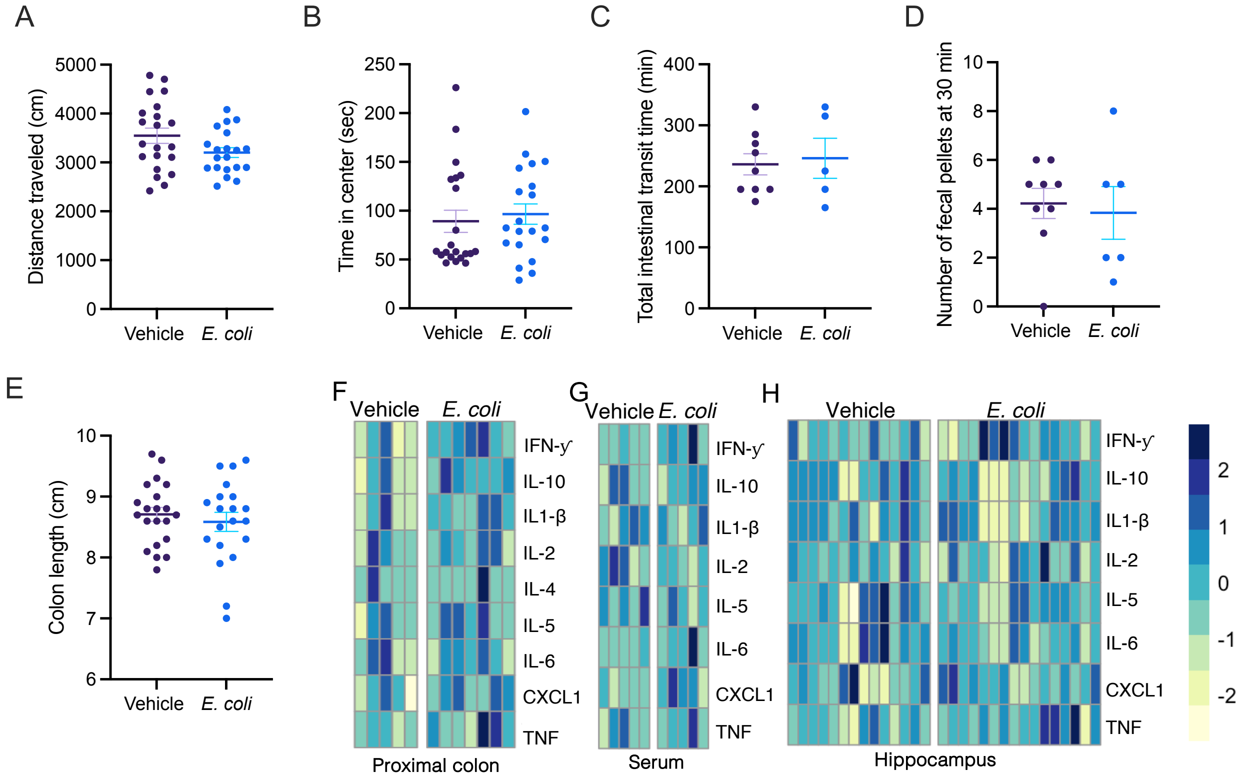


**Supplementary Figure S4, *associated with Fig. 6. E. coli* exposure does not induce sickness behavior or robust inflammation in 5xFAD mice.** Vehicle and *E. coli* exposed 5xFAD mice were tested for signs of sickness behavior on a battery of tests. **A)** Motor and **B)** anxiety-like behavior were measured on the open filed test. Gastrointestinal function was measured by **C)** carmine red elution and **D)** total fecal output during a 30-minute period. Inflammation was measured within the colon via **E)** colon length at the time of sacrifice and inflammatory cytokines and chemokines were measured within the **F)** proximal colon, **G)** serum, and **H)** hippocampus by multiplex ELISA. n = 5-21. Dots (or columns in **F-H**) represent individual mice, bars represent mean ± SEM. In **A-E** groups were compared using a two tailed t test. In **F-H,** groups were compared using multiple t tests adjusted for multiple comparisons using two-stage step-up (Benjamini, Krieger, and Yekutieli).

### Supplementary Figure S5.

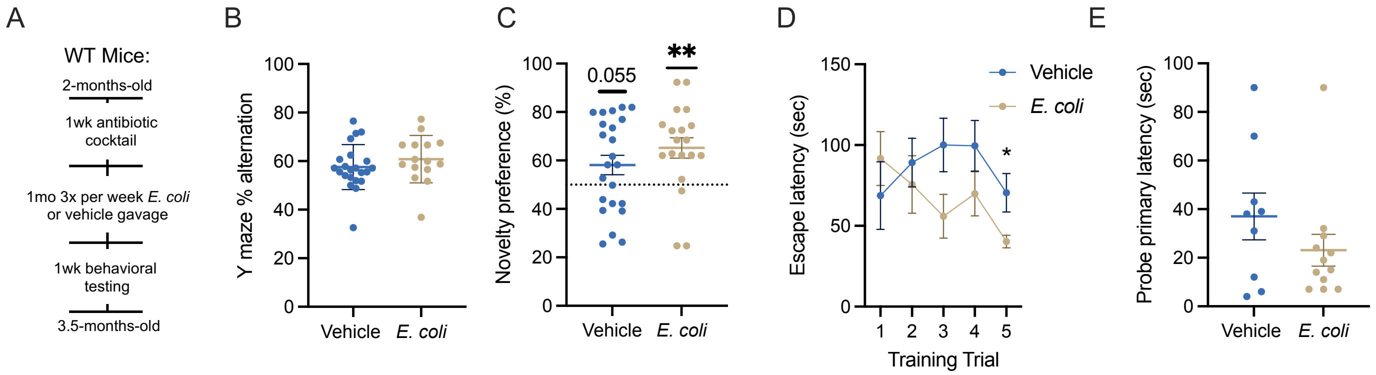


**Supplementary Figure S5, *associated with Fig. 6. E. coli* exposure does not induce cognitive impairment in wild type mice. A)** Wild type littermates were tested side by side with 5xFAD mice to see if *E. coli* exposure was sufficient to induce cognitive decline in the absence of familial AD mutations. Performance on the **B)** Y maze, **C)** object location test and **D-E)** Barnes maze. n = 9-23. Dots represent individual mice bars represent mean ± SEM. Groups were compared using a T test in **B** and **E**, one sample T test compared to 50% chance level in **C**, and two-way repeated measures ANOVA in **D**. *p < 0.05; ** p < 0.01.
